# Cooperative involvement of zipper-interacting protein kinase (ZIPK) and the dual-specificity cell-division cycle 14A phosphatase (CDC14A) in vascular smooth muscle cell migration

**DOI:** 10.1101/2024.03.06.583600

**Authors:** Abdulhameed Al-Ghabkari, David A. Carlson, Timothy A. J. Haystead, Justin A. MacDonald

## Abstract

Zipper-interacting protein kinase (ZIPK) is a Ser/Thr protein kinase with regulatory involvement in vascular smooth muscle cell (VSMC) actin polymerization and focal adhesion assembly dynamics. ZIPK silencing can induce cytoskeletal remodeling with disassembly of actin stress fiber networks and coincident loss of focal adhesion kinase (FAK)-pY397 phosphorylation. The link between ZIPK inhibition and FAK phosphorylation is unknown, and critical interactor(s) and regulator(s) are not yet defined. In this study, we further analyzed the ZIPK-FAK relationship in VSMCs. The application of HS38, a selective ZIPK inhibitor, to coronary artery vascular smooth muscle cells (CASMCs) suppressed cell migration, myosin light chain phosphorylation (pT18&pS19) and FAK-pY397 phosphorylation as well. This was associated with the translocation of cytoplasmic FAK to the nucleus. ZIPK inhibition with HS38 was consistently found to suppress the activation of FAK and attenuate the phosphorylation of other focal adhesion protein components (i.e., pCas130, paxillin, ERK). In addition, our study showed a decrease in human cell-division cycle 14A phosphatase (CDC14A) levels with ZIPK-siRNA treatment and increased CDC14A with transient transfection of ZIPK. Proximity ligation assays (PLA) revealed CDC14A localized with ZIPK and FAK. Silencing CDC14A showed an increase of FAK-pY397 phosphorylation. Ultimately, the data presented herein strongly support a regulatory mechanism of FAK in CASMCs by a ZIPK-CDC14A partnership; ZIPK may act as a key signal integrator to control CDC14A and FAK during VSMC migration.

## INTRODUCTION

Zipper-interacting protein kinase (ZIPK, also known as death-associated protein kinase 3, DAPK3 (Kawai et al. 1998; Kogel et al. 1998)) is a Ser/Thr protein kinase implicated in the regulatory control of a wide variety of cellular activities, including programmed cell death, autophagy, cell proliferation, migration and invasion, chromatin organization and remodeling, cleavage furrow ingression during cell division, and contractility. Sequence analysis of ZIPK revealed a high degree of sequence similarity (∼80%) for the N-terminal kinase domain within the DAPK family (Kogel et al. 1998). However, ZIPK does possess structural features that distinguish it from DAPK1 and DAPK2 (Chen and MacDonald 2023); specifically, the lack of a calmodulin-binding domain that enables signaling activation independent of changes in intracellular Ca^2+^ concentrations ([Ca^2+^]*_i_*) and the presence of a leucine zipper (LZ) domain that is important for homo- and hetero-dimerization events. Conceivably, these unique characteristics lead to divergent signaling function(s) and modulatory networks to control ZIPK activity (Haystead 2005; Gozuacik and Kimchi 2006; Bialik and Kimchi 2006; Shiloh, Bialik, and Kimchi 2014; Usui, Okada, and Yamawaki 2014).

ZIPK is known to regulate vascular smooth muscle cell (VSMC) motility and contraction through the diphosphorylation of the 20-kDa regulatory light chains of myosin II (LC20) at Ser19 and Thr18 (Murata-Hori et al. 1999; Niiro and Ikebe 2001; Borman et al. 2002; Komatsu and Ikebe 2004, 2014; MacDonald et al. 2016); phosphorylation of the myosin phosphatase-targeting subunit 1 (MYPT1) of myosin light chain phosphatase (MLCP) at Thr697 and Thr855 inhibitory sites (MacDonald, Borman, et al. 2001; Borman et al. 2002; Endo et al. 2004; Haystead 2005; Moffat et al. 2011; MacDonald et al. 2016); and the phosphorylation of the protein kinase C (PKC)– potentiated inhibitory protein for myosin phosphatase of 17 kDa (CPI-17) at Thr38 (MacDonald, Eto, et al. 2001). These targets are key mediators of the Ca^2+^ sensitization process in VSMCs (i.e., smooth muscle contraction in the absence of a change in [Ca^2+^]*_i_*).

Several reports implicate ZIPK in the regulation of cell morphology. The ectopic expression of the kinase in non-muscle cells contributes to membrane blebbing, cell rounding, and detachment from the cell matrix (Komatsu and Ikebe 2004; Murata-Hori et al. 1999). ZIPK can also contribute to the development and maintenance of myosin II-driven contractility and migration of non-muscle cells; studies show ZIPK to serve as a resistance mechanism against therapies targeted at melanoma (Orgaz et al. 2020) as well as a central participant in cytoskeletal reorganization and apoptotic processes associated with tumor cell survival, migration and epithelial-mesenchymal transformation in colon cancer (Chen and MacDonald 2021). In another example, an interaction of ZIPK with the prostate apoptosis response-4 (Par-4) protein facilitates accessibility to the cytoskeleton to influence VSMC contractility (Vetterkind and Morgan 2009; Vetterkind et al. 2010). Additional reports detail the impact of ZIPK on actin cytoskeleton networks and focal adhesion assembly dynamics (Nehru, Almeida, and Aspenstrom 2013; Zheng, Gong, and Zhen 2020; Turner et al. 2023).

The cell adhesion and motility of VSMCs are regulated by the transduction of external stimuli to the intracellular milieu via transmembrane sensors (i.e., integrins) that drive multiple tyrosine phosphorylation events and assembly of focal adhesion protein complexes (Goldmann 2012). Integral to this complex process, FAK phosphorylation at Tyr397 creates a motif that is recognized by various Src homology 2 (SH2) domain-containing proteins which, in turn, leads to the conformational activation of Src and stimulation of cytoskeletal changes (Kleinschmidt and Schlaepfer 2017; Mitra, Hanson, and Schlaepfer 2005). The tension development that occurs during VSMC contraction is promoted by dynamic actions on the actin cytoskeleton (Ohanian, Pieri, and Ohanian 2015; Tang and Gerlach 2017). Indeed, the formation of the actin filament network at submembranous locations enhances membrane rigidity and facilitates the transmission of tension generated by cross-bridge cycling events associated with contractile activity (Gunst and Zhang 2008; Opazo Saez et al. 2004; Zhang and Gunst 2008). Intriguingly, ZIPK over-expression in human fibroblast cells was observed to alter actin filament assembly and induce the reorganization focal adhesion components with a coincident suppression of FAK (Tyr397) phosphorylation status (Nehru, Almeida, and Aspenstrom 2013). A recent study from our group has further demonstrated that suppression of ZIPK signaling with a small-molecule inhibitor or siRNA-mediated knockdown resulted in pronounced alterations in cell morphology as well as cytoskeletal architecture of VSMCs with coincident reduction in focal adhesion kinase (FAK) phosphorylation (Turner et al. 2023). Nonetheless, the precise mechanism through which ZIPK regulates FAK phosphorylation and the focal adhesion complex remains undefined.

Dual-specificity protein phosphatases (DUSPs) have emerged as key regulators of cell migration and adhesion events (Larsen, Tremblay, and Yamada 2003). In particular, the human cell-division cycle 14A (CDC14A) phosphatase regulates the cytoskeletal organization and is co-localized with F-actin at the leading cell edges of migratory cells (Chen et al. 2017; Chen et al. 2016). CDC14A knockout promotes enhanced cell motility and is associated with a decrease in cell adhesion with actin cytoskeleton rearrangement and dephosphorylation of key regulators, including the epithelial protein lost in neoplasm (EPLIN, also known as LIMA1) and extracellular signal-regulated kinase (ERK). The intracellular phosphoprotein substrates of CDC14A and their associated protein kinases are only now being identified (Chen et al. 2017); however, emerging evidence supports a functional relationship between CDC14A and ZIPK (Ye et al. 2014; Wu et al. 2015). Further examination of the ZIPK-CDC14A relationship is required to understand its mechanistic consequence on actin cytoskeleton focal adhesion organization.

In this study, we develop novel mechanistic understanding of a ZIPK and FAK signaling nexus that also integrates the human CDC14A phosphatase. Central to our experiments is the application of a validated ZIPK small molecule inhibitor, arylthiopyrazolo[3,4-d]pyrimidinone (HS38) that competitively inhibits ZIPK (Carlson et al. 2013; Carlson et al. 2018). HS38 is a cell-permeable, potent ZIPK inhibitor with few of the off-target liabilities associated with other small molecule DAPK inhibitors (Al-Ghabkari et al. 2016; MacDonald et al. 2016; Carlson et al. 2013); thus, HS38 provides a unique opportunity to examine ZIPK-dependent effects on the focal adhesion complex and the actin cytoskeleton, as well as the involvement of CDC14A therein.

## MATERIALS and METHODS

### Materials

All chemicals were of reagent grade or higher, and unless otherwise stated were purchased from BioRad, Sigma-Aldrich or VWR. AlexaFluor488-Phalloidin, AlexaFluor568 goat anti-rabbit IgG, AlexaFluor568 goat anti-mouse IgG and AlexaFluor488 goat anti-rabbit IgG were purchased from ThermoFisher Scientific (Mississauga, ON, Canada). Antibodies were from Cell Signaling Technology (Danvers, MA: anti-β-actin, 4967S; anti-ERK, 9102S; anti-pERK(pT202/pY204), 4370S; anti-tubulin, 2144A; anti-p130Cas(pY249), 4014A; anti-p130Cas(pY410), 4011S; anti-p130Cas(E1L9H), 13856; and anti-LC20(pST18/pS19), 3764; anti-Src(pY527), 2105S), Santa Cruz Biotechnology (Dallas, TX: anti-Lamin B, sc-6216; anti-LC20, sc-15370), and Abcam (Cambridge, UK: anti-ZIPK/ZIPK, ab151956; anti-FAK, ab40794; anti-FAK(pY397), ab39967; anti-FAK(pY576), ab55335; anti-paxillin(pY118), ab4833; anti-SM22α, ab14106; anti-CDC14A, ab10536). Human ZIPK-siRNA (am-4390824) and scrambled siRNA (am-AM4613) were purchased from Ambion/Thermofisher Scientific, and human CDC14A siRNA and scrambled siRNA (i504211) were from Applied Biological Materials (Richmond, BC, Canada). FAK-14 inhibitor was from Tocris Bioscience (Bristol, UK). ZIPK inhibitor HS38 was synthesized as previously described) (Carlson et al. 2013). Phos-tag acrylamide was from Wako Chemicals (Richmond, VA). The Duolink *In Situ* Proximity Ligation Assay kit was purchased from Olink Bioscience (Uppsala, Sweden).

### Vascular smooth muscle cell cultures

Coronary artery smooth muscle cells (CASMCs, CC-2583; Lonza; Allendale, NJ) were maintained and cultured in smooth muscle basal media (SmGM-2 BulletKit; CC-3182; Lonza). The media was supplemented with hEGF, insulin, hFGF-B and FBS. Cells were confirmed to be negative for *Mycoplama* strain contamination and then seeded at density of 3,500 viable cells/cm^2^ according to the manufacturer’s protocol. Cell viability, morphology and proliferative capacity were routinely examined after recovery from cryopreservation. Passages from 8-12 were used in different experimental conditions. In some cases, CASMCs were transiently transfected with Flag-tagged ZIPK. Cells were maintained in SmGM-2 media (Lonza) supplemented with 10% (v/v) FBS (Invitrogen/Thermofisher Scientific). DNA vectors were transfected using FuGENE 6 transfection reagent (Roche Applied Science, Laval, QC) according to the manufacturer’s instructions. Briefly, a transfection mixture of Opti-MEM (200 μL; Invitrogen) and FuGENE reagent (10 μL) was incubated for 5 min with subsequent addition of 5 μg DNA and a further 20 min incubation. The transfection mixture was then added to confluent (70-80%) CASMCs which were maintained at 37 °C for 20-48 h. Whole cell extracts were prepared from CASMC cultures by first washing cells with PBS (136.9 mM NaCl, 2.7 mM KCl, 10.1 mM Na_2_HPO_4_ and 1.76 mM KH_2_PO_4_) and then completing cellular lysis with addition of 2x sample buffer (2% (w/v) SDS, 100 mM DTT, 10% (v/v) glycerol, and 60 mM Tris-HCl, pH 6.8) with gentle rocking followed by heating at 95 °C.

### ZIPK knockdown with siRNA

CASMCs (2 × 10^5^) were seeded in 2 ml antibiotic-free normal growth medium supplemented with FBS. Scrambled- and ZIPK-siRNAs were diluted with siRNA transfection medium (sc-36868, Santa Cruz Biotechnology) to give a final concentration of 20 nM. The siRNA solutions were gently overlaid onto the cells and incubated for 6-12 h at 37 °C. Normal growth medium (2 ml containing 2 times the normal serum and antibiotic concentrations) was added, and the cells were screened for 24-48h until harvest for analyses.

### Co-immunoprecipitation

Cells were washed 3x with cold PBS prior to lysis with 150 mM NaCl, 1% (v/v) NP-40, 2 mM EDTA, 1 mM DTT, 1 mM PMSF, 1mM benzamidine, Complete protease inhibitor cocktail (Roche) and 50 mM Tris-HCl, pH 7.4. Cellular lysates were cleared by centrifugation (14,000 rpm, 20 min) and then incubated (30 min) with Protein G-Sepharose beads (GE Healthcare, Mississauga, ON, Canada) followed by subsequent centrifugation (14,000 rpm, 10 min). Supernatants were removed for immunoprecipitation with mouse monoclonal anti-CDC14A antibody (overnight at 5 °C). To capture the antigen-antibody complex, Protein G-Sepharose beads were incubated with supernatant mixture for 4 h with rocking followed by centrifugation (14,000 rpm for 10 sec). Supernatant was discarded and beads were washed 3x with ice-cold PBS.

### Western blot analyses

Cell lysates were resolved by SDS-PAGE and transferred to 0.2 μm nitrocellulose membranes in a Tris/glycine transfer buffer containing 10% (v/v) methanol. In some cases, Phos-tag SDS–PAGE was used to examine protein phosphorylation profiles. Proteins were transferred to polyvinylidene difluoride (PVDF) membranes at 25 V for 16 h at 4 °C. Proteins were fixed on the membrane with 0.5% (v/v) glutaraldehyde in PBS for 45 min and then washed with TBST (25 mM Tris-HCl, 137 mM NaCl, 3 mM KCl, and 0.05% (v/v) Tween-20). Nonspecific-binding sites were blocked with 5% (w/v) nonfat dry milk in TBST. Membranes were washed with TBST and incubated overnight with primary antibody at a 1:1,000 dilution in 1% (w/v) nonfat dry milk in TBST. Membranes were incubated for 1 h with horseradish peroxidase (HRP)-conjugated secondary antibody (dilution 1:10,000) and developed with enhanced chemiluminescence (ECL) reagent (GE Healthcare). All western blots were visualized with a LAS4000 Imaging Station (GE Healthcare), ensuring that the representative signal occurred in the linear range. Quantification was performed by densitometry with ImageQuant TL software (GE Healthcare).

### Wound healing assay

An *in vitro* wound-healing or “scratch” assay was performed to measure unidirectional migration of CASMCs. Cells were seeded at 1 × 10^5^ cells/well and then incubated for 48 h at 37 °C in a humidified atmosphere of 5% CO_2_. Once the cells reached 70%-80% confluency, vehicle (DMSO) and small molecule ZIPK or FAK inhibitor were applied to the cells and then incubated for 30 min followed by washing twice with PBS. A wound area was carefully created for cell migration by scraping the cell monolayer with a sterile pipette tip. Detached cells were gently rinsed away with PBS and an image taken by phase-contrast microscopy. Subsequently, another set of images were taken after 24 h time interval. The extent of migration was estimated by quantifying the migration area into the cell free region.

### Immunocytochemistry

CASMCs (2 × 10^5^ cells) were plated in SmGM-2 media with 10% (v/v) FBS at 37 °C with 5% CO2. Cells were fixed (15 min) in 4% (v/v) paraformaldehyde in PBS and then permeabilized with 0.5% (v/v) Tween-20 for 10 min. Cells were then incubated at 4 °C overnight with primary antibody diluted 1:100 in blocking serum (0.3% BSA, 5% goat serum, 0.3%, Triton X-100 in PBS, pH 7.4). After washing with PBS, immunoreactivity was detected with goat anti-rabbit or mouse anti-rabbit AlexaFluor488-conjugated secondary antibody. For F-actin visualization, AlexaFluor488-phalloidin was diluted 1:40 in 1% (w/v) BSA in PBS and then incubated with the cells for 1 h at room temperature. Cells were rinsed with PBS, counterstained with DAPI for 5 min to detect nuclei, and then examined with an InCell 6000 Imaging System (GE Healthcare). For visualization, the InCell 6000 Imaging System was programmed to complete whole-well scanning of four independent plates of cells. Ten random visual fields were analyzed from each well, and eight images were taken from each field. The F-actin fluorescence signal was quantified with ImageJ (https://imagej.nih.gov) by splitting the color channels of the original image. The AlexaFluor488-phalloidin signal was identified by selecting cells of interest using the drawing tool and then applying the analysis parameters of the area and integrated density functions. Immunofluorescence signal was quantified and then normalized to the DAPI nuclear counterstain.

### Proximity ligation assay

CASMCs were utilized for proximity ligation assays (PLA) using Duolink *in situ* PLA detection kit (Olink). Cells were fixed (15 min) in 4% (v/v) paraformaldehyde in PBS and then permeabilized with 0.5% Tween-20 for 10 min. Cells were then washed with PBS, blocked for 30 min at 37 °C in Duolink blocking solution, and incubated overnight at 4 °C with pairs of primary antibodies in Duolink antibody diluent solution. Cells were washed 3x with PBS and then were incubated with secondary antibodies conjugated with Duolink PLA PLUS and MINUS probes for 2 h at 37 °C. Anti-mouse PLUS and anti-rabbit MINUS were employed to detect complexes between ZIPK-CDC14A, FAK-CDC14A and FAK(pY397)-CDC14A. The hybridization signal was amplified using amplification solution containing DNA polymerase enzyme and then detected using red fluorescent signal using the InCell 6000 Imaging System. Proximity signals are represented as red dots and quantified using NIH ImageJ. Control experiments employed only one primary antibody for each protein.

### Data analysis

Data are presented as the mean ± SEM, with n indicating the number of independent experiments. Significance was assessed with the Student’s *t*-test with P < 0.5 considered to indicate statistical significance. All statistical analyses were performed using the GraphPad Prism v6.0 software.

## RESULTS

### ZIPK can be effectively targeted in CASMCs with siRNA or small molecule HS38 inhibitor

ZIPK expression was effectively suppressed in CASMCs with an optimized siRNA treatment protocol (Figure 1A). The link between ZIPK and myosin regulatory light chain (LC20) diphosphorylation at T18 and S19 has been extensively described in different cell and tissue models (Murata-Hori et al. 1999; Niiro and Ikebe 2001; Borman et al. 2002; Komatsu and Ikebe 2004; Carlson et al. 2013; Komatsu and Ikebe 2014; MacDonald et al. 2016), and we identified a corresponding decrease in LC20-2P (pT18/pS19) levels in CASMCs treated with siRNA-ZIPK or HS38 inhibitor when compared to the respective control condition (Figure 1B). Immunocytochemistry examination for the dual phosphorylation of LC20 (pT18/pS19) using phospho-specific antibody showed that the total LC20-2P signal was significantly attenuated in CASMCs treated with HS38 compared to vehicle control (Figure 1C). Furthermore, the distribution of LC20-2P was altered such that the immunoreactivity was no longer uniformly aligned with actomyosin filaments but was found more randomly distributed throughout the cell in punctate aggregates. In Figure 1D, immunostaining of CASMCs for ZIPK revealed a prominent nuclear pool with an additional diffuse signal observed throughout the cytoplasm. Finally, the application of HS38 to CASMCs also resulted in attenuated cell migration during scratch-wound healing (Figure 1E).

**FIGURE 1.**
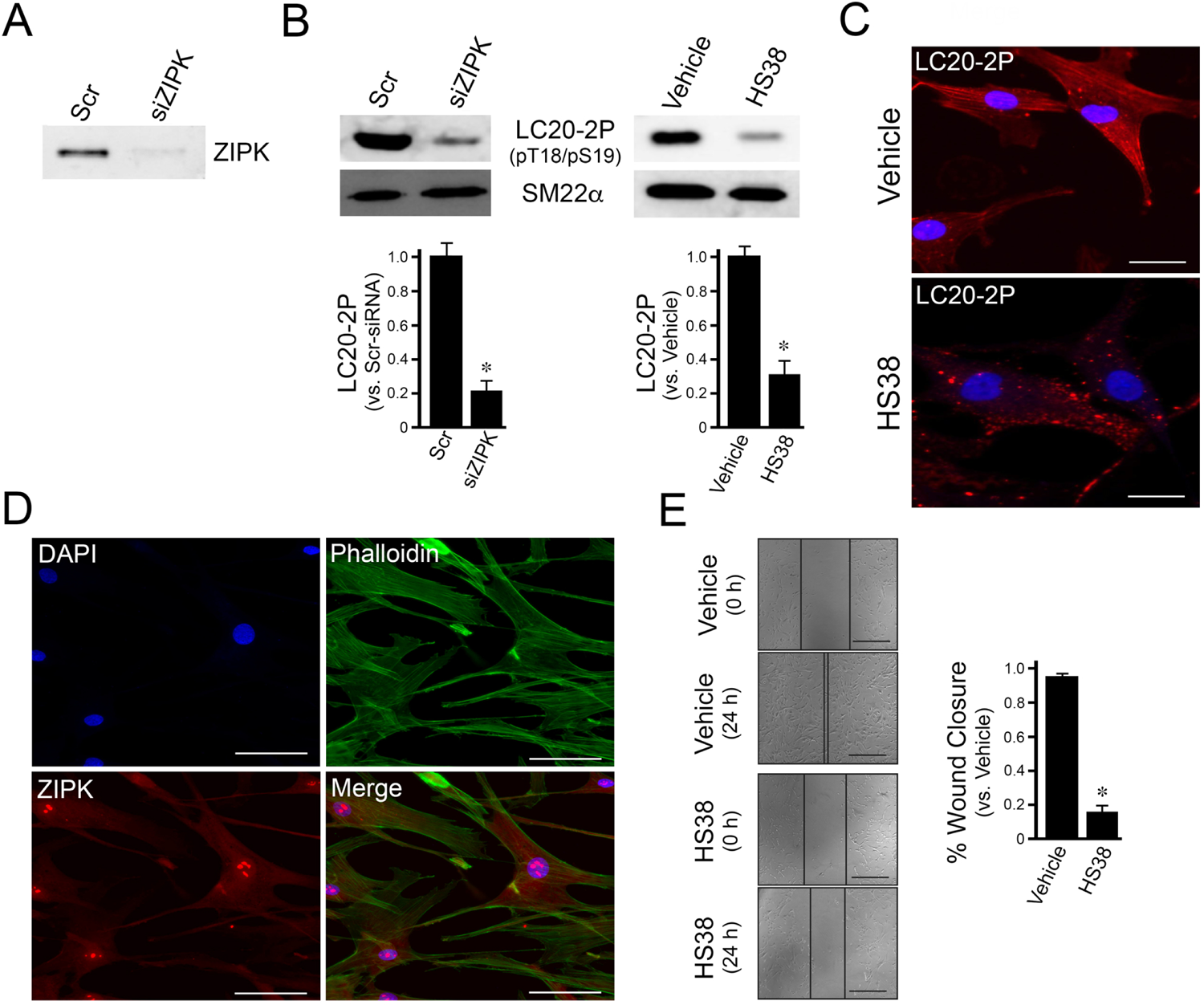
ZIPK inhibition by HS38 impact LC20 phosphorylation in CASMCs. CASMCs were interrogated by ZIPK silencing with siRNA-mediated knockdown or pharmacological inhibition. (*A*) ZIPK-siRNA (siZIPK) or scrambled control siRNA (Scr) was applied to CASMCs, then ZIPK protein was examined by western blotting and densitometric quantification of total cell lysates. (*B*) The phosphorylation status of ZIPK substrate LC20 (diphosphorylation of pT18/pS19) was analyzed by western blotting of total cell lysates from CASMCs treated with siRNA (siZIPK or Scr) and small molecule inhibitor (HS38, 25 μM or DMSO vehicle control). SM22α was used as a loading control. (*C*) Subsequent to vehicle or HS38 treatments, CASMCs were fixed and reacted with anti-LC20-2P(pT18,pS19), AlexaFluor568-conjugated secondary antibodies (red) and DAPI nuclear stain (blue) before preparation for confocal immunofluorescence microscopy. (*D*) CASMCs were fixed and stained with AlexaFluor488-phalloidin (green), anti-ZIPK with AlexaFluor568-conjugated secondary antibody (red) and DAPI nuclear stain (blue) to examine ZIPK subcellular localization. Scale bars = 30 μm. (*E*) Confluent CASMCs were serum-starved (24 h) in the presence of HS38 (25 μM) and subjected to a wound healing scratch assay. Cell migration was monitored, and wound closure was expressed as the % coverage of the initial cell-free zone. Values represent means ± S.E.M. for n=4 independent experiments. *Significantly different from the vehicle control (Student’s *t*-test, p < 0.05).

### Changes in cytoskeletal structure and FAK phosphorylation follow HS38 treatment of CASMCs

CASMCs treated with HS38 or siRNA-ZIPK demonstrate a significant disassembly of central stress fibers causing cell shrinkage and shape change (Turner et al. 2023). Stress fibres and cortical actin are continuously (de)stabilized by focal adhesion kinase (FAK)-regulated processes (Mitra, Hanson, and Schlaepfer 2005). Treatment with HS38 was previously shown to reduce FAK-pY397 phosphorylation (Turner et al. 2023). These findings were confirmed since concentration-dependent effects of HS38 were also observed on FAK-pY397 phosphorylation (Figure 2A). Examination of FAK-pY397 phosphorylation was also completed using immunocytochemical analysis of CASMCs treated with HS38. In this case, CASMCs showed an attenuation in total signal intensity for FAK-pY397 with HS38 treatment (Figure 2B).

**FIGURE 2.**
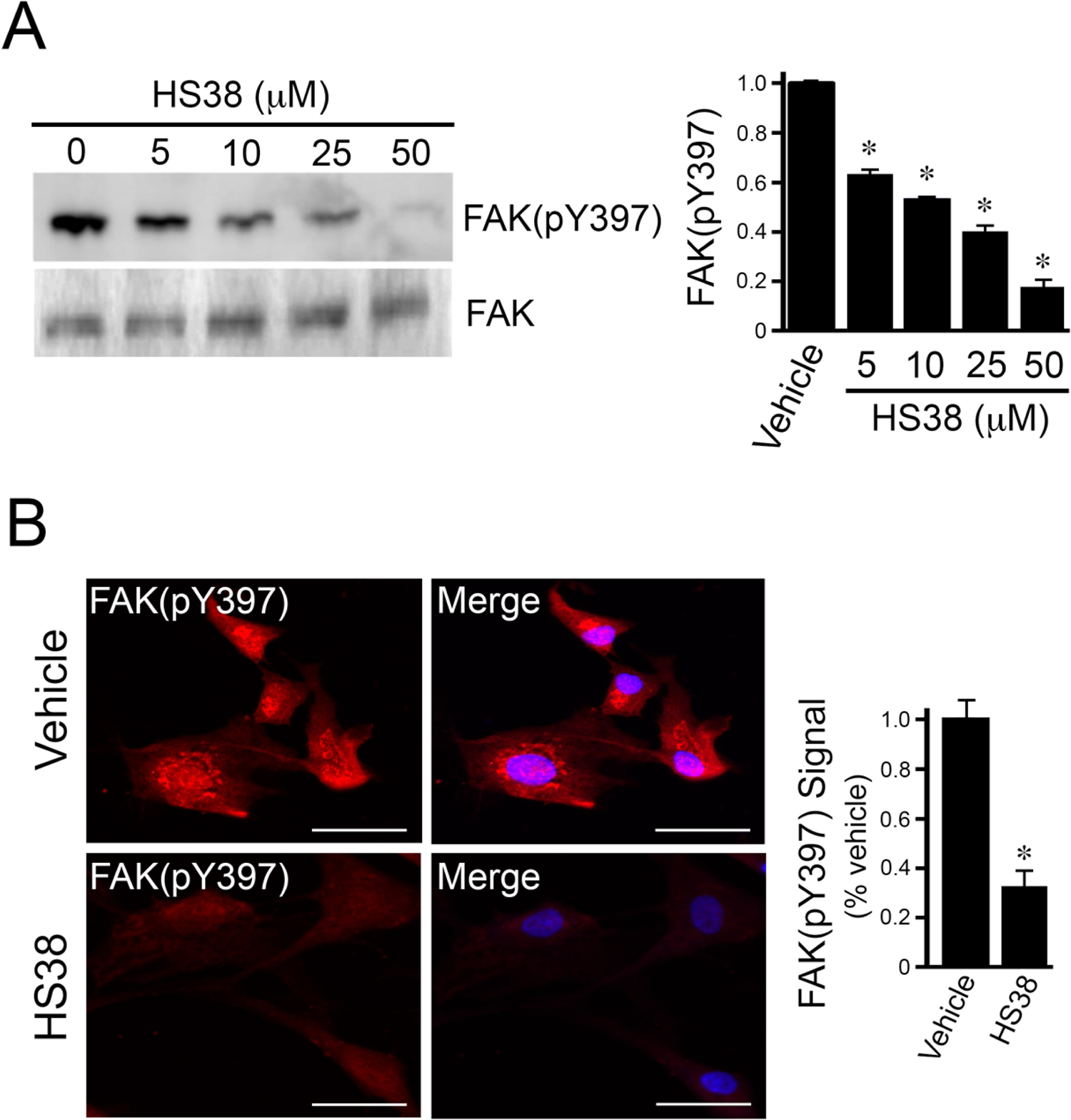
Effect of HS38 treatment on cytoskeletal architecture and FAK phosphorylation of vascular smooth muscle cells. (*A*) CASMCs were treated with increasing [HS38] for 16 hr, and FAK-pY397 phosphorylation was quantified with normalization to total FAK load. (*B*) Immunocytochemistry was completed on CASMCs fixed and reacted with anti-[pY397]-FAK, AlexaFluor568-conjugated secondary antibodies (red) and DAPI nuclear stain (blue) before preparation for confocal immunofluorescence microscopy. Total cellular immunofluorescence signal intensity for FAK-pY397 was quantified. \**P* < 0.05 versus vehicle controls; Student’s *t*-test, n=5 independent experiments. Scale bars = 30 μm.

### HS38 treatment of CASMCs results in differential phosphorylation of various smooth muscle and focal adhesion proteins

Given the impact of ZIPK inhibition on FAK signaling events, we also investigated the effect of HS38 treatment on various proteins involved in the focal adhesion complex. We examined the inhibitory phosphorylation site (pY527) of the proto-oncogene tyrosine-protein kinase (Src) and the docking protein p130Cas (Crk-associated substrate) phosphorylation sites (pY249 and pY410). No significant changes in the phosphorylation of Src-pY527 or p130Cas-pY249 were observed (Figure 3A); however, a significant decrease in p130Cas-pY410 phosphorylation was detected (Figure 3B). Paxillin (pY118) phosphorylation and ERK activation events are also linked to focal adhesion assembly. Application of HS38 to CASMCs reduced the total immunofluorescence signal obtained with a paxillin(pY118)-specific antibody (Figure 3C). A significant decrease of paxillin-pY118 was quantified by western blot analysis when CASMCs were treated with either siRNA-ZIPK or HS38 compound (Figure 3D). The activating phosphorylation of ERK (pT202/pY204) was also suppressed following the application of increasing concentrations of HS38 inhibitor to CASMCs (Figure 3E).

**FIGURE 3.**
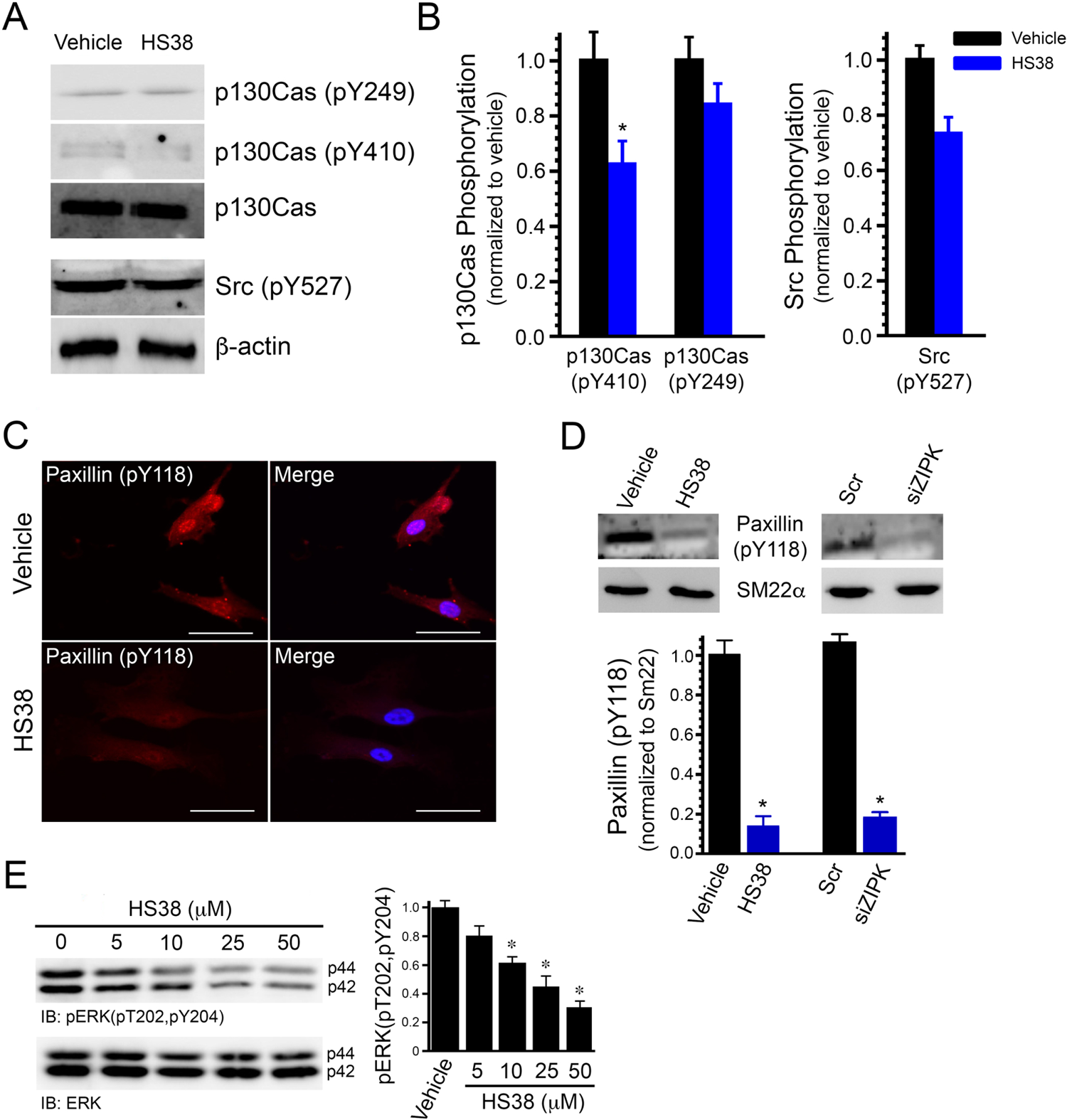
Effect of HS38 treatment on the phosphorylation of focal adhesion proteins in CASMCs. (*A-B)* Proteins from whole cell lysates of CASMCs incubated for 16 h with HS38 (25 μM) or DMSO (vehicle) were examined by western blotting and densitometric quantification following immunoreaction with anti-[pY249]-p130Cas, anti-[pY410]-p130Cas and anti-[pY527]-Src. β-actin and p130Cas(E1L9H) and were used as a loading controls. \**P* < 0.05 versus vehicle controls; Student’s *t*-test, n=3 independent experiments. (*C*) Immunocytochemistry was completed on CASMCs fixed and reacted with anti-[pY118]-paxillin and AlexaFluor568-conjugated secondary antibodies (red) or DAPI nuclear stain (blue) before preparation for confocal immunofluorescence microscopy. Scale bars = 30 μm. (*E*) The phosphorylation of paxillin (pY118) following HS38 or siRNA-ZIPK treatment was quantified by western blotting with SM22α muscle protein used as loading control. (*F*) CASMCs were treated with increasing [HS38] for 16 hr. Whole cell lysates were prepared, and phospho-ERK(pT202,pY204) was quantified by western blotting with normalization to total ERK protein. \**P* < 0.05 versus vehicle controls; Student’s t-test, n=7 independent experiments.

### Intracellular localization of CDC14A with ZIPK and FAK in CASMCs

Dual specificity phosphatase human cell-division cycle 14A (CDC14A) is known to be associated with the actin cytoskeleton of human cells (Chen et al. 2017; Chen et al. 2016). Following immunocytochemistry analyses of CASMCs, CDC14A immunoreactivity was localized to phalloidin-stained F-actin filaments under basal conditions (Figure 4A, panel i). As shown previously in Figure 1D, ZIPK immunostaining under basal conditions revealed a prominent nuclear pool with an additional diffuse signal observed throughout the cytoplasm (Figure 4B, panel i). The overlap of immunofluorescence signals suggests that CDC14A and ZIPK are generally localized in similar areas of the CASMC (Figure 4B, panel i). The application of HS38 to CASMCs altered the intracellular localization of CDC14A; the immunoreactivity was no longer associated with F-actin filaments and was spread diffusely throughout the cytoplasm (Figure 4A, panel ii). Diffuse ZIPK immunoreactivity was still found in the cytoplasm following HS38 treatment; however, there was a marked decrease in the nuclear signal observed (Figure 4B, panel ii). Further analyses of CASMCs were characterized by prominent co-localization of CDC14A and FAK immunofluorescence intensities under control conditions (Figure 4C, panel i). The co-localization of CDC14A and FAK immunofluorescence signals was also profoundly reduced following HS38 treatment of CASMCs (Figure 4C, panel ii).

**FIGURE 4.**
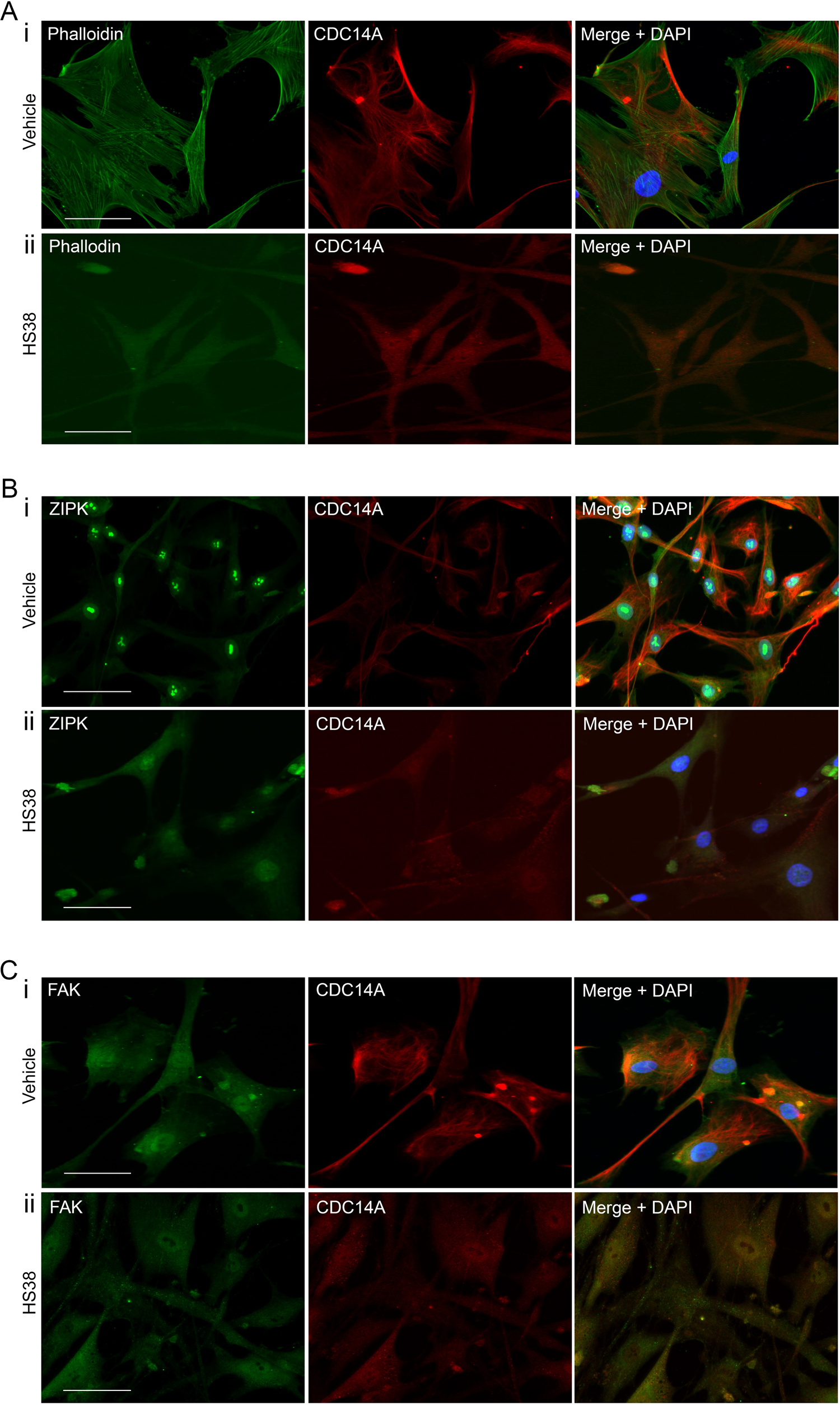
HS38 treatment results in rapid dephosphorylation and nuclear accumulation of FAK. (*A*) Time-dependent FAK-pY397 dephosphorylation was examined in CASMCs treated with HS38 (25 μM) for 0 – 30 min. (*B*) Alterations in total FAK phosphorylation with HS38 treatment (0 – 50 μM) were examined by Phos-tag SDS-PAGE and western blotting with anti-FAK antibody. (*C* and *D*) CASMCs were fixed and reacted with rabbit anti-[pY397]-FAK or anti-FAK in the presence of AlexaFluor488-conjugated secondary antibodies (green) and with DAPI nuclear stain (blue) before preparation for confocal immunofluorescence microscopy. Representative images from 4 independent experiments are shown for cells treated in the absence (*C*; DMSO, vehicle control) or presence of HS38 (*D*; 25 μM, 10 min). Scale bars = 30 μm. (*E*) Nuclear and cytoplasmic fractions were prepared and analyzed by western blotting for FAK, tubulin (cytoplasmic marker), and lamin B (nuclear marker).

We further investigated the putative CDC14A, ZIPK and FAK signalosome using the proximity-ligation assay (PLA). In this case, fluorescent dots (i.e., PLA events) are observed *in situ* when the targets of two primary antibodies are localized in close proximity (∼30 nM (Soderberg et al. 2006)). As shown in Figure 5A, a noteworthy number of PLA events for ZIPK and CDC14A were detected in CASMCs. Likewise, significant numbers of PLA events were recorded for CDC14A and FAK (Figure 5B). Controls were completed using antibodies for each protein alone to assess the specificity of the assay and to eliminate false positives. Taken together, the immunofluorescence microscopy and PLA data are strongly supportive of an integrated ZIPK-CDC14A-FAK signalosome in CASMCs.

**FIGURE 5.**
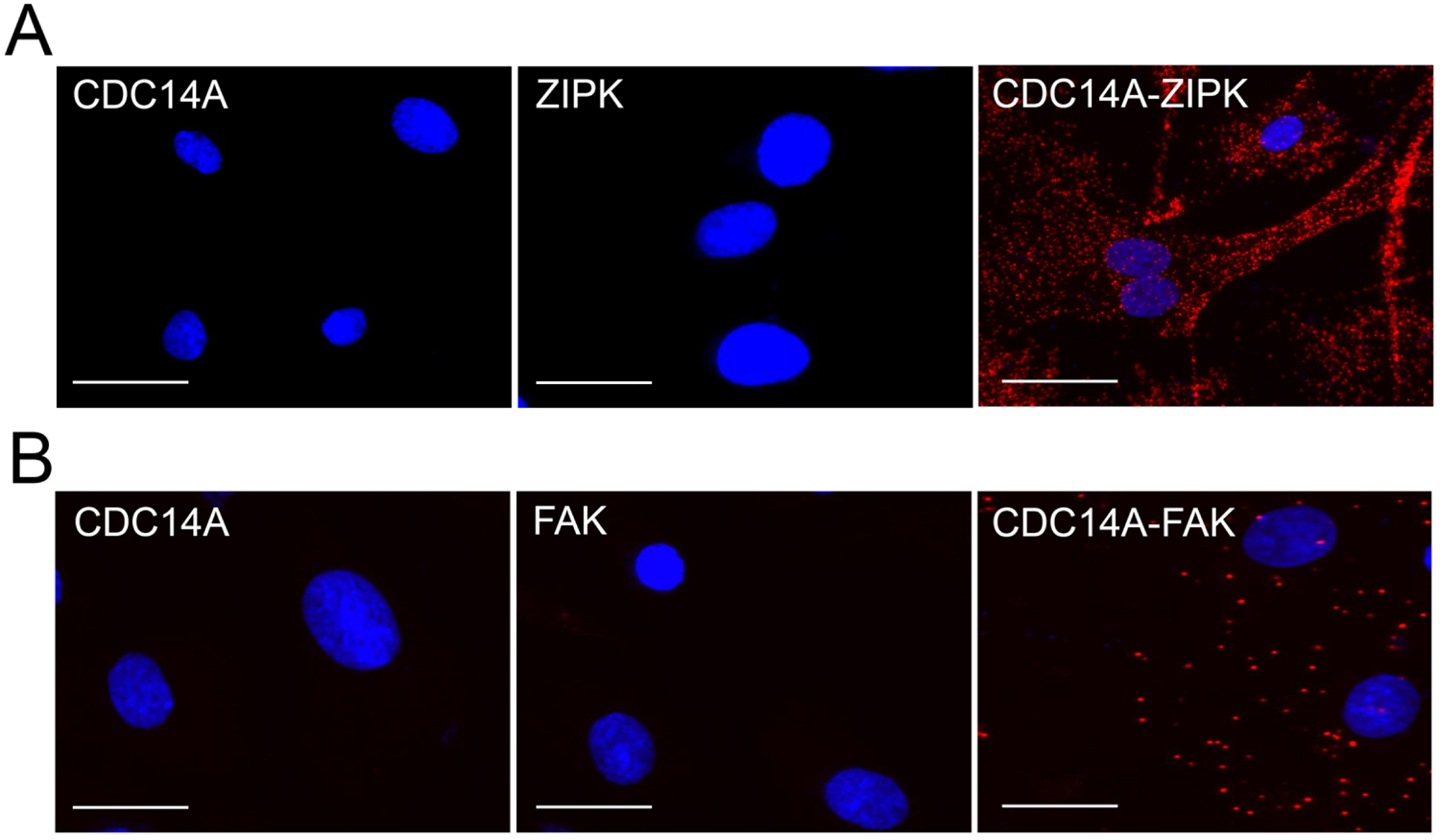
Effects of HS38 treatment on the co-localization of dual-specificity phosphatase CDC14A with F-actin filaments, ZIPK and FAK in CASMCs. After 16 h incubation with vehicle (DMSO) or small molecule inhibitor (HS38, 50 μM), CASMCs were fixed and prepared for confocal immunofluorescence microscopy of endogenous proteins. (*A*) CASMCs were incubated with AlexaFluor488-phalloidin to examine F-actin cytoskeletal organization (green), or anti-CDC14A and AlexaFluor568-conjugated secondary antibodies to examine CDC14A (red) following treatment with vehicle (*panel i*) or HS38 (*panel ii*). (*B*) CASMCs were incubated with anti-ZIPK and AlexaFluor488-conjugated secondary antibodies (green), or anti-CDC14A and AlexaFluor568-conjugated secondary antibodies (red) following treatment with vehicle (*panel i*) or HS38 (*panel ii*). (*C*) CASMCs were incubated with anti-FAK and AlexaFluor488-conjugated secondary antibodies (green), or anti-CDC14A and AlexaFluor568-conjugated secondary antibodies (red) following treatment with vehicle (*panel i*) or HS38 (*panel ii*). Representative images from 3 independent experiments are shown with DAPI nuclear stain (blue).

### Regulation of CDC14A by ZIPK in CASMCs

As analyzed by immunofluorescence microscopy, treatment of CASMCs with HS38 disrupted the association of CDC14A with the cytoskeleton (Figure 4). Moreover, qualitative differences in the co-localization of CDC14A with ZIPK and FAK signals were also apparent (Figures 4 & 5). So, additional experiments were designed to interrogate the putative ZIPK-CDC14A partnership. The over-expression of Flag-ZIPK in CASMCs was associated with a ∼4-fold increase in CDC14A (Figure 6A). The silencing of ZIPK by siRNA-knockdown was associated with a marked loss of CDC14A protein (Figure 6B), as detected by western blotting. Co-immunoprecipitation of ZIPK was possible with anti-CDC14A antibody (Figure 6C). But, the co-immunoprecipitation of ZIPK from lysates was not detected following the treatment of CASMCs with HS38. These data indicate that the amount of CDC14A protein in CASMCs is directly associated with ZIPK levels and imply that HS38-binding by ZIPK could disrupt the interaction. In a final set of experiments, the impact of HS38 on the ZIPK-CDC14A-FAK signalosome was examined with PLA (Figure 7A). HS38 treatment was demonstrated to markedly diminish the number of ZIPK-CDC14A proximity events in CASMCs (Figure 7B), a result that parallels the immunoprecipitation findings shown previously in Figure 6C. Further to this, the number of PLA events recorded for CDC14A and FAK as well as CDC14A and FAK-pY397 were greatly reduced in CASMCs treated with HS38 (Figure 7B).

**FIGURE 6.**
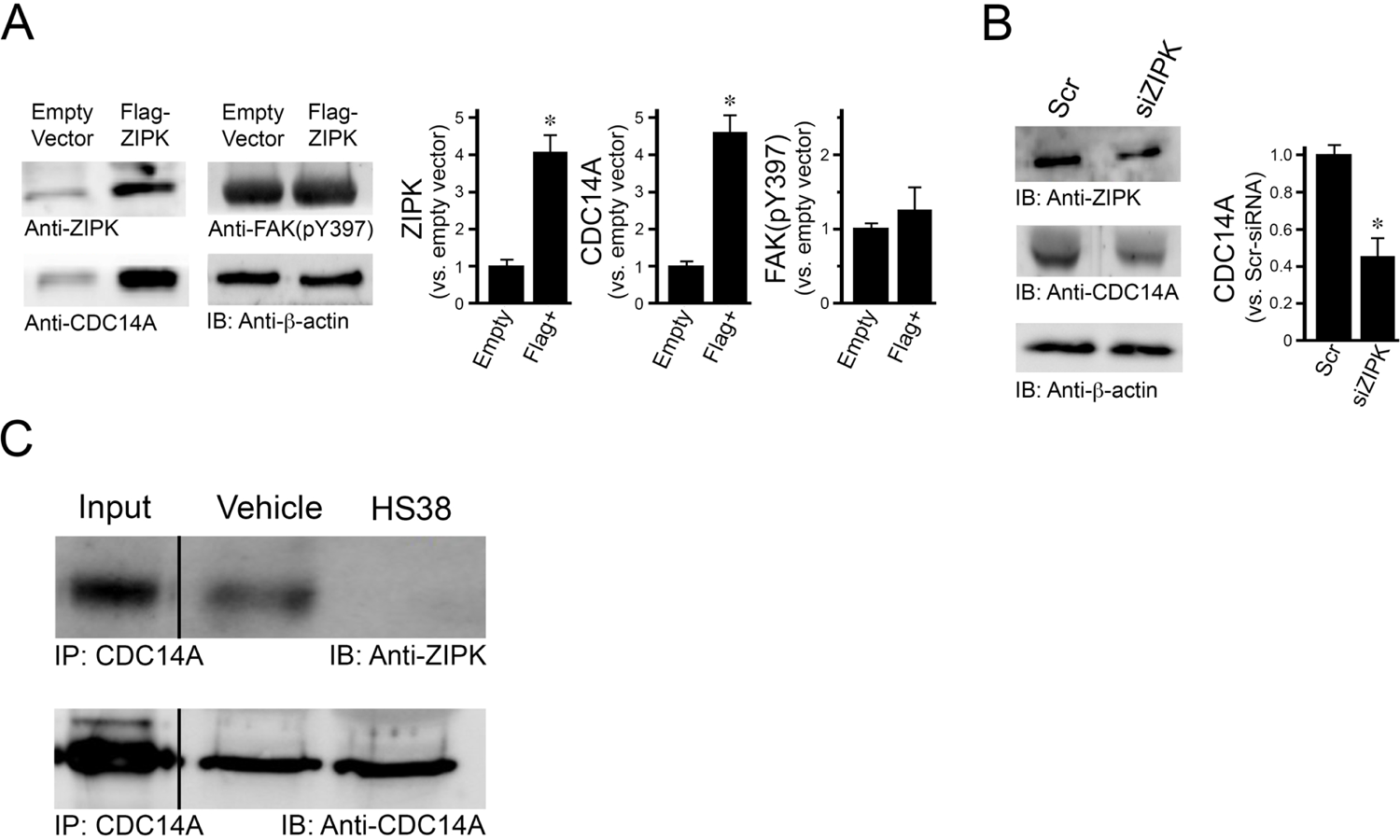
Impact of ZIPK expression on CDC14A and FAK in CASMCs. Alterations in ZIPK, CDC14A and FAK(pY397) were examined by western blotting and densitometric quantification in whole cell lysates following manipulation of ZIPK levels. (*A*) CDC14A levels and FAK-pY397 phosphorylation status were examined in CASMCs subjected to over-expression of FLAG-tagged ZIPK. (*B*) Following treatment with siRNA-ZIPK or control Scr-siRNA, whole cell lysates were prepared and analyzed for CDC14A. (*C*) Immunoprecipitation was performed on CASMC extracts with anti-CDC14A antibody. Samples of the total cell lysate (input protein) as well as content bound to Protein G-Sepharose were analyzed by SDS-PAGE and western blotting with antibodies against ZIPK and CDC14A proteins. Blots are representative of three independent experiments. β-actin was used as loading control in all cases. \**P* < 0.05 versus controls; Student’s *t*-test, n=4 independent experiments.

**FIGURE 7.**
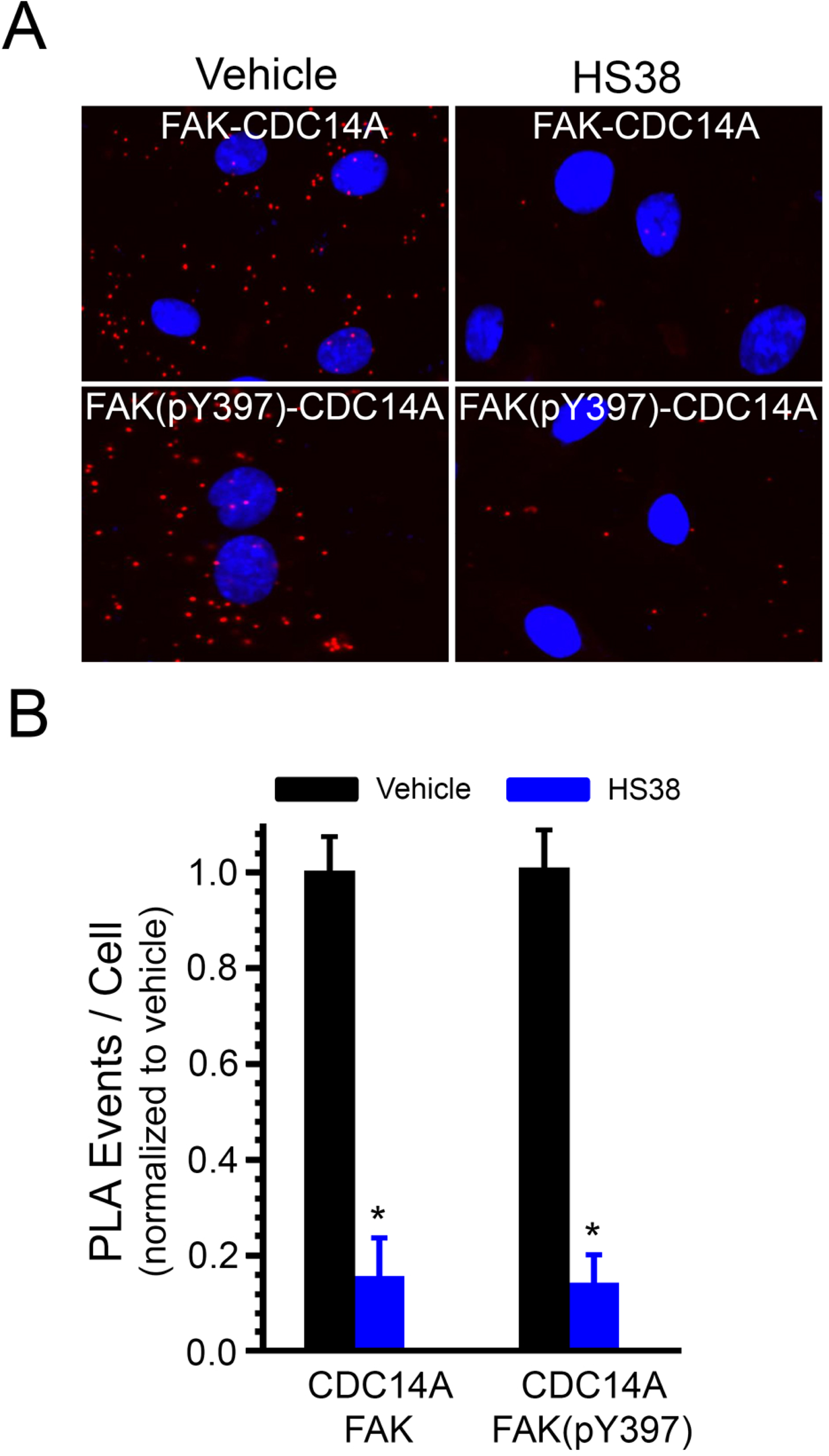
HS38 treatment reduces proximity events associated with CDC14A and FAK(pY397) signaling complex formation in CASMCs. (*A*) The proximity-ligation assay (PLA) was used to examine CDC14A/FAK and CDC14A/FAK-pY397 co-localization events after treatment with vehicle (DMSO) or HS38 (25 μM). CASMCs were reacted with primary antibody for CDC14A in the presence or absence of anti-FAK or anti-FAK-pY397. (*B*) PLA events (red dots) for CDC14A/ZIPK, CDC14A/FAK and CDC14A/FAK-pY397 were identified by confocal microscopy and quantified following administration of vehicle (DMSO) or HS38. Scale bars = 30 μm. \**P* < 0.05 versus vehicle controls; Student’s t-test, n=4 independent experiments.

## DISCUSSION

Previously, we reported that administration of HS38 to human CASMCs could induce cytoskeletal remodeling with disassembly of intermediate filaments and central stress fiber networks (Turner et al. 2023). The phenotypic changes were associated with significant loss of FAK-pY397 phosphorylation. Now, we confirm the impact of ZIPK inhibition with HS38 on the FAK phosphorylation status using independent methods (i.e., PLA and co-localization of immunofluorescence) and also identify HS38 treatment to attenuate the activity of important focal adhesion members (i.e., pCas130, paxillin) and downstream modulators (i.e., ERK). Cell microscopy of PLA events showed ZIPK to be in close proximity to the dual phosphatase CDC14A and activated FAK-pY397; these localization events diminished with HS38 exposure. Importantly, we reveal a critical partnership for CDC14A and ZIPK since our results showed decreased CDC14A protein with ZIPK-siRNA treatment and increased CDC14A protein with ectopic expression of ZIPK. Co-immunoprecipitation data suggests that HS38 could block the interaction of ZIPK with CDC14A. The data presented herein strongly support a regulatory mechanism of FAK and focal adhesion dynamics by a ZIPK-CDC14A partnership. Ultimately, ZIPK may act as a key signal integrator to control CDC14A and the FAK-dependent regulation of cellular adhesion events.

Several new pieces of evidence support the impact of ZIPK on cytoskeleton-related cellular functions. ZIPK-dependent effects on FAK signaling were previously observed in fibroblasts where WT-ZIPK overexpression resulted in the reorganization of stress fibers with increased focal adhesion size and marked decrease in FAK-pY397 (but not FAK-pY576) signal (Nehru, Almeida, and Aspenstrom 2013). Moreover, the ectopic expression of kinase-dead ZIPK^D161A^ did not impact upon the focal adhesion size but was still observed to suppress FAK-pY397. Our findings were distinct from those observed by Nehru and coauthors. Namely, the overexpression of WT-ZIPK in CASMCs had no effect on FAK-pTyr397 levels. Both inhibition of ZIPK with HS38 and silencing of ZIPK with siRNA-mediated knockdown elicited cytoskeletal remodeling with loss of FAK-pY397 phosphorylation signal (Turner et al. 2023). ZIPK inhibition with HS38 caused focal adhesion disassembly and induced the translocation of FAK to the nucleus. The over-expression of a ZIPK mutant lacking the leucine zipper (ZIPKΛ1LZ) was previously shown to have no effect on FAK-pY397 levels (Nehru, Almeida, and Aspenstrom 2013); however, our knockdown of ZIPK with siRNA did elicit cytoskeletal remodeling with loss of pY397 phosphorylation signal (Turner et al. 2023). The unique cellular environments (i.e., fibroblasts vs. VSMCs) may account for the different findings. Ultimately, more research is needed to identify whether scaffolding and/or kinetic properties of ZIPK are required for its impact upon FAK activity and focal adhesion assembly in VSMCs. Our findings also indicate that FAK-pY576 phosphorylation was not altered by HS38 treatment of CASMCs. This phosphorylation is mediated by Src on the activation loop of the kinase domain to achieve maximal FAK activity, so data from both studies suggest that the kinase activity of FAK was not altered by ZIPK. Rather it is the assembly of protein complexes at the focal adhesion could be influenced. The fact HS38 had no impact on FAK-pY576 also supports a lack of off-target effect by HS38 on Src; likewise, past experiments that examined the off-target effects of HS38 did not identify any inhibitory potential toward FAK or Src in kinase activity screens.

The nuclear translocation of FAK was identified when CASMCs were treated with the ZIPK inhibitor HS38. Although this is a novel report for ZIPK influence on FAK cellular localization, several other publications have previously reported changes in the intracellular localization of FAK. The nuclear translocation of FAK was previously detected in the hypertrophic myocardium of spontaneously hypertensive heart failure rats, which is suggestive of a role for FAK in the regulation of mechanical transduction through modulation of gene transcription in cardiac myocytes (Peng et al. 2006). The nuclear shuttling of FAK was also reported in a variety of cellular functions, including the regulation of cell survival by promoting turnover of p53 (tumor suppressor) activity through the murine double minute-2 (Mdm2)-dependent p53 ubiquitination pathway (Lim et al. 2008), downregulation of VCAM-1 expression by promoting GATA4 transcription factor ubiquitination (Lim et al. 2012), and heterochromatin remodeling by the transcriptional induction of CpG-binding protein 2 (MBD2) through activation of histone deacetylase 1 (HDAC1) dissociation (Luo et al. 2009). Similar observations were reported when human umbilical vein endothelial cells (HUVECs) were treated with staurosporine (a broad-acting kinase inhibitor) to induce FAK nuclear localization that was accompanied by FAK dephosphorylation and loss of focal adhesion contacts (Lobo and Zachary 2000).

The localization of CDC14A in CASMCs was mainly confined to the F-actin cytoskeleton. The co-localization of immunofluorescence signals suggested an association between ZIPK and CDC14A, mainly in the cytoplasmic space that includes the cytoskeleton. PLA experiments further confirmed the spatial overlap of CDC14A and ZIPK proteins, their association with F-actin stress fibers, and proximity to FAK. Wu and colleagues have previously demonstrated a direct interaction of ZIPK and CDC14A proteins using yeast 2-hybrid screening, *in vitro* pull-down assays, and co-immunoprecipitation (Wu et al. 2015). The authors hypothesized that ZIPK(ZIPK) could stimulate CDC14A phosphatase activity by direct phosphorylation although this putative mechanism was not validated with subsequent experimentation. It is tempting to speculate that CDC14A acts as a FAK phosphatase to target the pY397 site. Although CDC14A is classified as a dual specificity phosphatase, phospho-proteomic investigations by Chen and colleagues show CDC14A to act preferentially on pSer residues (Chen et al. 2017); thus, it is unlikely that CDC14A directly targets the pY397 site of FAK.

Intriguingly, we also show an upregulation of CDC14A protein following ZIPK overexpression; a decrease in CDC14A protein was also detected with siRNA-mediated knockdown of ZIPK. A similar relationship for CDC14A and ZIPK was previously communicated by Ye and colleagues using high glucose-stimulated human aortic smooth muscle cells (Ye et al. 2014). Here, shRNA-silencing of ZIPK provided concurrent attenuation of CDC14A levels. Conversely, ectopic expression of ZIPK was associated with upregulation of CDC14A in this aortic VSMC. The CDC14A-ZIPK partnership is further evidenced by the RNA sequencing profiles of 9,736 tumors and 8,587 normal samples deposited in the TCGA and the GTEx databases. We examined the gene expression of *ZIPK* and *CDC14A* in various tumor types using the GEPIA pipeline (Figure 8), and the analyses revealed higher *ZIPK* transcript levels with increased *CDC14A* expression in two types of cancers when compared to a normal dataset: pancreatic adenocarcinoma (PAAD) and thymoma (THYM). This indicates a potential expression co-dependency of *ZIPK* and *CDC14A*.

**FIGURE 8.**
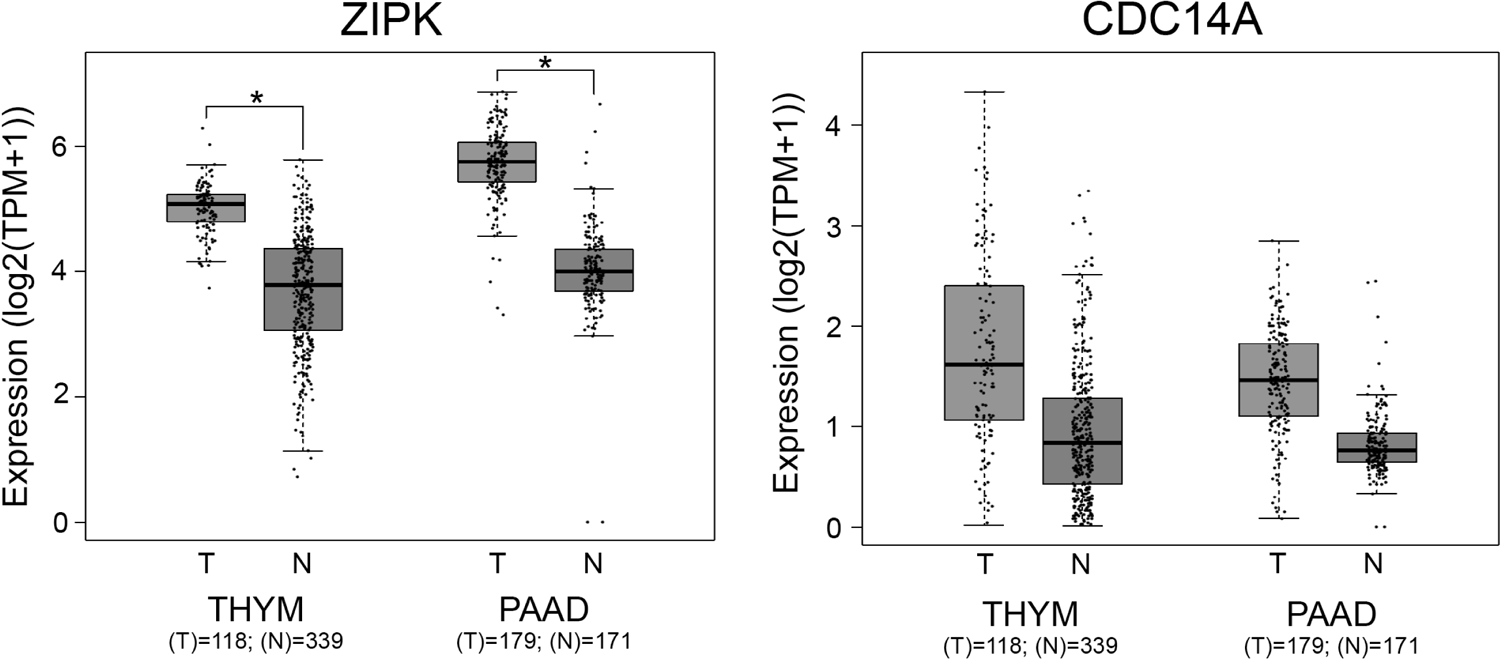
Expression analysis of *ZIPK* and *CDC14A* in tumors using the GEPIA pipeline. Differential expression analyses were completed for thymoma (THYM) and pancreatic adenocarcinoma (PAAD). Significant differences in differential expression (“TCGA tumors vs TCGA normal + GTEx normal” or “TCGA tumors vs TCGA normal”) were identified for the disease state (Tumor (T) or Normal (N)) using one-way ANOVA. \**P* < 0.05. The gene expression levels (y-axis) were log2(TPM+1) transformed for differential analysis with the log2FC defined as the median (“Tumor” - “Normal”).

Taken together, our findings support a unique partnership of ZIPK, CDC14A and FAK in CASMCs. The actions of CDC14A appear to impact on VSMC migration and FAK-dependent adhesion events. ZIPK is suggested to bind and mask CDC14A from degradation signals that allow the phosphatase to regulate FAK phosphorylation and focal adhesion events. The mechanism whereby ZIPK acts to titrate CDC14A levels remains undefined; however, Chen and colleagues report several regulators of ubiquitin- and SUMO-linked protein proteolysis in their BioID survey of the CDC14A interaction network (Chen et al. 2017). Future research should further interrogate the co-dependency of ZIPK and CDC14A with prioritization on the novel ability of ZIPK to suppress degradation mechanisms acting on CDC14A. Finally, it will be necessary to assess CDC14A phosphatase activity in CASMCs to evaluate its ability to control FAK phosphorylation and cell migration.

## ACKNOWLEDGEMENTS

This work was supported by a grant from the Canadian Institutes of Health Research (MOP#97931 to JAM) and the National Institutes of Health (R01DK065954-05 to TAJH). JAM held an Alberta Innovates – Health Solutions (AIHS) Senior Scholar Award and was recipient of a Canada Research Chair (Tier 2) in Smooth Muscle Pathophysiology.

## AUTHOR CONTRIBUTIONS

All persons designated as authors qualify for authorship, and all those who qualify for authorship are listed. A.A.-G. conducted the experiments, completed data analysis, prepared figures and co-wrote the manuscript. D.A.C. synthesized the HS38 compounds. T.A.J.H. coordinated the production of HS38 inhibitor and made intellectual contributions to the project. J.A.M. conceived and coordinated the study, assisted with experimental design, co-wrote the manuscript, provided trainee supervision, and made intellectual contributions to the project. All authors approved the final version of the manuscript and agree to be accountable for all aspects of the work in ensuring that questions related to the accuracy or integrity of any part of the work are appropriately investigated and resolved.

